# Spatial model of water quality in Southeast Asian urban lake based on HBI using macrozoobenthos diversity as a proxy

**DOI:** 10.1101/2021.02.16.431487

**Authors:** Andri Wibowo

## Abstract

Urban lake is one of ecosystem that has experienced anthropogenic pressures and this can affect its water quality. One of a robust approach to assess the water quality is by using Hilsenhoff Biotic Index (HBI). This tool is quite versatile since it can be applied by using any aquatic organism as proxy including macrozoobenthos. This invertebrate group also has an advantage since it is common and easy to collect. Here this study is first, aiming to provide HBI based water quality spatial model using macrozoobenthos as a proxy applied in urban lake in West Java in Southeast Asia and second to seek the best model that can represent the water quality variables in particular dissolved oxygen (DO), pH, and temperature. Based on the spatial model and HBI, either inlet or outlet parts of the lake, it has better water quality in comparison to central parts. Based on HBI values, water quality in inlet and outlet parts (HBI = 6.7) is categorized as fair and poor (HBI = 6.9) for the central parts of the lake. The increase in HBI and decrease in water quality are positively correlated with the increase in water temperature variable in comparison to water DO and pH variables. Akaike model selection confirms that the macrozoobenthos diversity can be used as a proxy for increase in water temperature (Ψ)_HBI_ (~temp)(AIC = −10.264) followed by combination of water temperature increase and decrease in DO(Ψ)_HBI_ (~temp+DO)(AIC = −9.042398).

## INTRODUCTION

Macrozoobenthos is aquatic animals that sensitive to natural and anthropogenic disturbances due to their low mobility and regarding to this condition, macrozoobenthos has been used widely as indicator to assess the health of aquatic environment and water quality. Many water quality related indices have been developed using macrozoobenthos as the proxy and this fall into one of three categories including diversity indices, similarity indices, or biotic index (Simić & Simić 1999, Zhongjun et al 2015). The macroinvertebrate biotic index is widely applied in developed countries and had been consequently optimized and standardized (Gabriels et al 2010), while in East Asia, its application is slowly but gradually growing [2, 9]. The successful use of macroinvertebrate indices abroad to track the health of water bodies and the number of extensive related studies in shallow lakes can be observed in these following studies (Gong & Xie 2001, Jiang et al. 2006).

Usng macrozoobenthos as proxy to assess water quality has several advantages. It has prompted resurgence in the use of biological assessment tools to monitor and evaluate aquatic systems (Karr & Dudley 1981) rather than simply measuring the chemical contents of surface water considering macrozoobenthos can represent the realt time relation between water quality and health of an aquatic ecosystem. As a result, for nearly a century macrozoobenthos have been used as a biological indicator of pollution. Macrozoobenthos are also particularly useful in modern rapid bio-assessments since they are having numerous advantages in comparison to other organisms. Macrozoobenthos is usually present in some form, numerous, and it has a relatively large size (Lenat et al. 1981). Macrozoobenthos is comparatively easy to identify in the field and macrozoobenthos life cycles are also often long enough to help measuring chronic environmental threats.

Several existing abiotic index or a score system including the Trent Biotic Index (TBI) or its offshoots (i.e. Extended Biotic Index, Chandler’s Score), New Zealand Macroinvertebrate Community Index (MCI) (Stark 1998), the Australia Stream Invertebrate Grade Number-Average Level(SIGNAL), the Wisconsin Biotic Index (BI) (Hilsenhoff 1987), the Biotic Monitoring Patagonian Stream (BMPS), and Hilsenhoff Biotic Index (HBI) were found to be effective because of their sensitivity and objectivity. HBI has been used widely to assess the water qualtiy. Whereas which specific water quality variables are best represented by the HBI are still poorl understood. Most of HBI studies were also lack of spatial interpretation while spatial distribution of HBI is required and very improtant to locate precisley the water bodies with poor water qualtiy and determine the pollution sources.

## MATERIALS AND METHODS

### Study area

This studied lake was a lake located in the middle of city. This urban lake was geographically located in populated Depok City in West Java Province. The lake has inlet in its south part and outlet in the north part. The lake is surrounded by vegetation and secondary forests. Whereas, the central part in the east side of the lake is closed to the road and anthropogenic activities. Macrozoobenthos was sampled from inlet in south and outlet in north parts of the lake and central parts.

### Method

#### HBI

The Hilsenhoff Biotic Index (HBI) estimates the overall tolerance of the macrozoobenthos community in a sampled area by weighing the relative abundance using taxonomic group including family and genus as a proxy. Macrozoobenthos are assigned a tolerance number from 0 to 10 pertaining to that organism's known sensitivity to organic pollutants; 0 being most sensitive, 10 being most tolerant. The HBI in each sampling station is described as following equation:

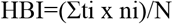

With: ti = tolerance index of family i, ni = number of individual of family i, N = total number of individual of all species. The values of tolerance index of each macrozoobenthos were obtained from literatures as can be seen in Table 1 below.

**Table 1.** Tolerance index of several macrozoobenthos

| Family | $ti$ | References |
| --- | --- | --- |
| Lymneaidae | 6 | Djumanto et al. (2013), Rachman et al (2016), Badri (2020) |
| Pachychilidae | 8 | Rustiasih (2018) |
| Planorbidae | 7 | Wijeyaratne & Kalaotuwawe (2017), Kahirun et al. (2019) |
| Thiaridae | 6 | Wijeyaratne & Kalaotuwawe (2017), Rustiasih (2018) |
| Viviparidae | 6 | Kahirun et al. (2019) |

The HBI values were classified into several classes with the lowest values were categorized as good and the highest values were categorized as poor. The water quality class based on HBI values can be seen in Table 2 below.

**Table 2.** HBI and water quality class

| Value ranges | Water quality |
| --- | --- |
| 0.0-3.60 | No pollution |
| 3.70-4.60 | Very good |
| 4.70-5.60 | good |
| 5.70-6.60 | Fair |
| 6.70-7.60 | poor |
| 7.70-8.60 | Very poor |
| 8.70-10.00 | No organism |

#### Spatial model

Spatial model used in this paper is addressed to assess the HBI distribution values in lake. The HBI spatial model was developed using interpolation method. The obtained HBI values were tabulated into GIS table and estimated using inverse distance weight method.

#### Akaike HBI model selection

HBI model as function of water quality variables including water dissolved oxygen (DO), pH, and temperature was developed using Akaike Information Criterion (AIC). The AIC was developed using the linear regression. The measured parameters included in AIC are R^2^ and adjusted R^2^. To build the model, 3 explanatory covariates including water DO, pH, and temperature and combinations of those covariates were included in the analysis to develop the model.

## RESULTS AND DISCUSSION

### Spatial model of HBI

The Figure 2 presents the spatial distribution of HBI values and water quality across the urban lake. The HBI values range from 6.7 to 6.9 and the water quality in urban lake is categorized as fair and poor. From the spatial model, it is very clear that the higher HBI values (6.9) were mostly distributed in the central parts of the lake. Whereas the lower HBI values of 6.7 were mostly distributed in the inlet in south parts and outlet in north parts. Meaning to say that the water quality in the central parts of the lake was lower than north and south parts of the lake. The range of water quality variables including water DO, pH, and temperature was available in Figure 3. The water was ranged from 29.5 °C to 30.5 °C, pH was ranged from 6.0 to 6.4, and range of 4.5 – 6.0 mg/l for DO (Figure 3).

**Figure 1.**
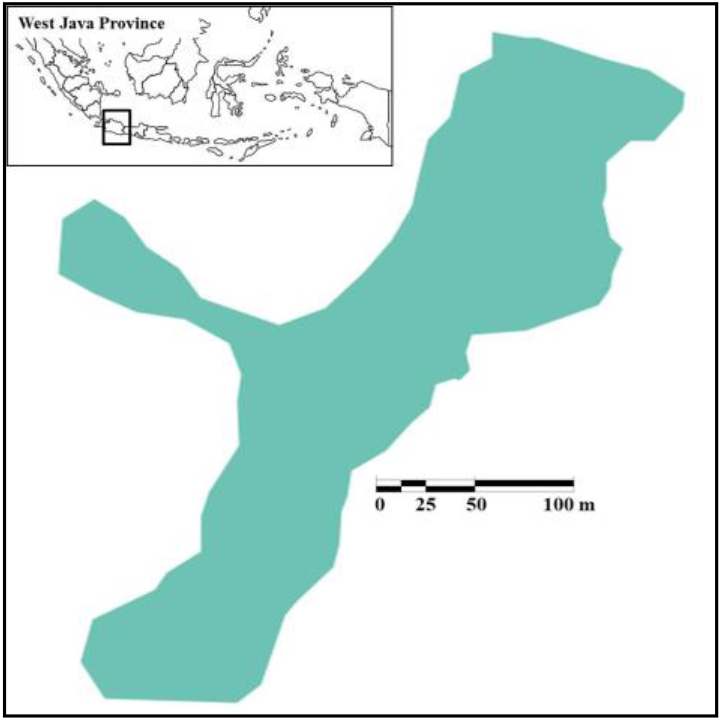
Location of study area in an urban lake in West Java Province, Indonesia.

**Figure 2.**
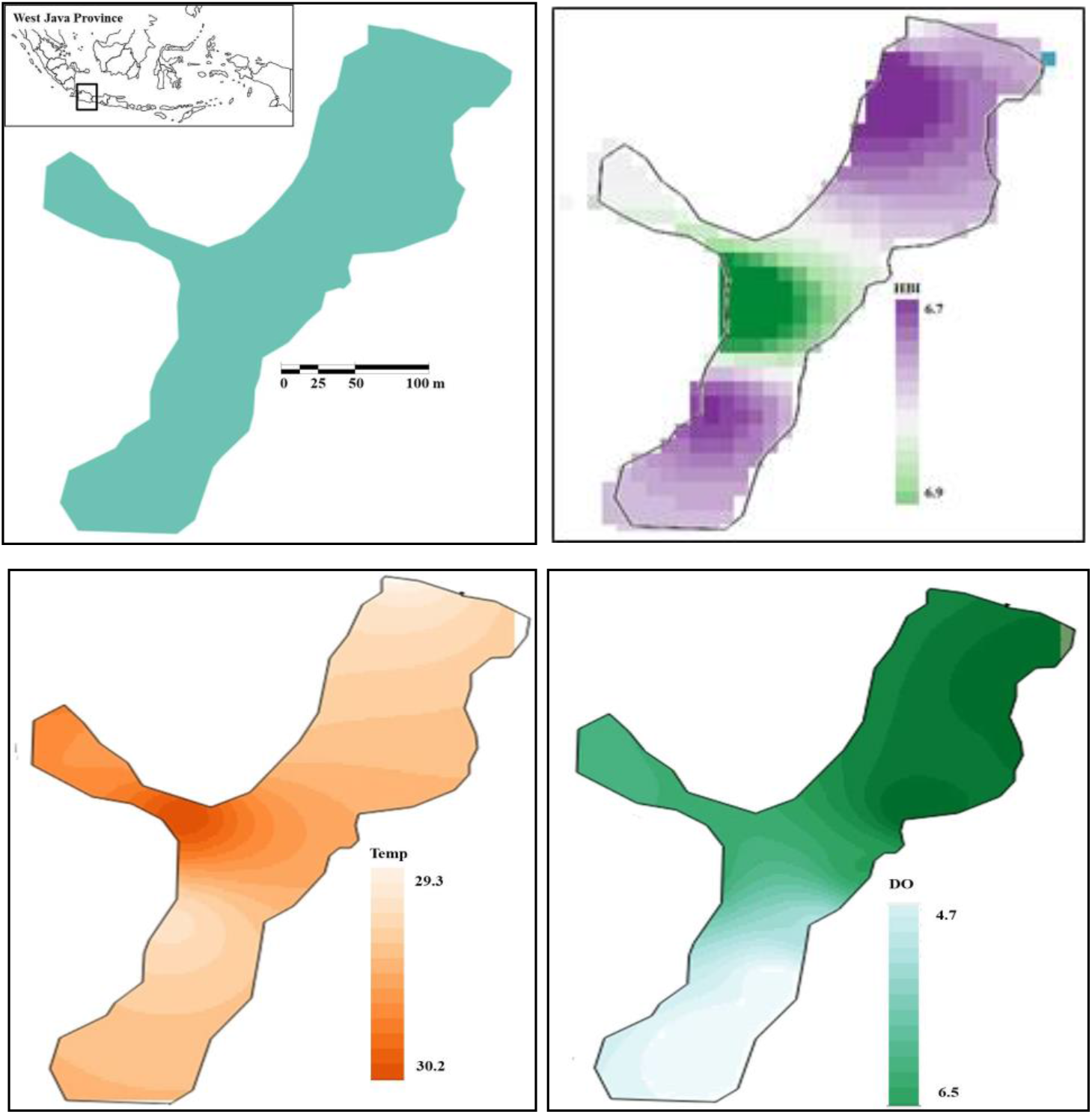
First row: studied urban lake (left) and spatial model (right) of HBI values (6.7-6.9) in studied urban lake with high values (6.9) and poor water quality observed in the central parts of the lake. Second row: spatial model (left) of temperature (°C) and DO (mg/l) (right) in studied urban lake.

**Figure 3.**
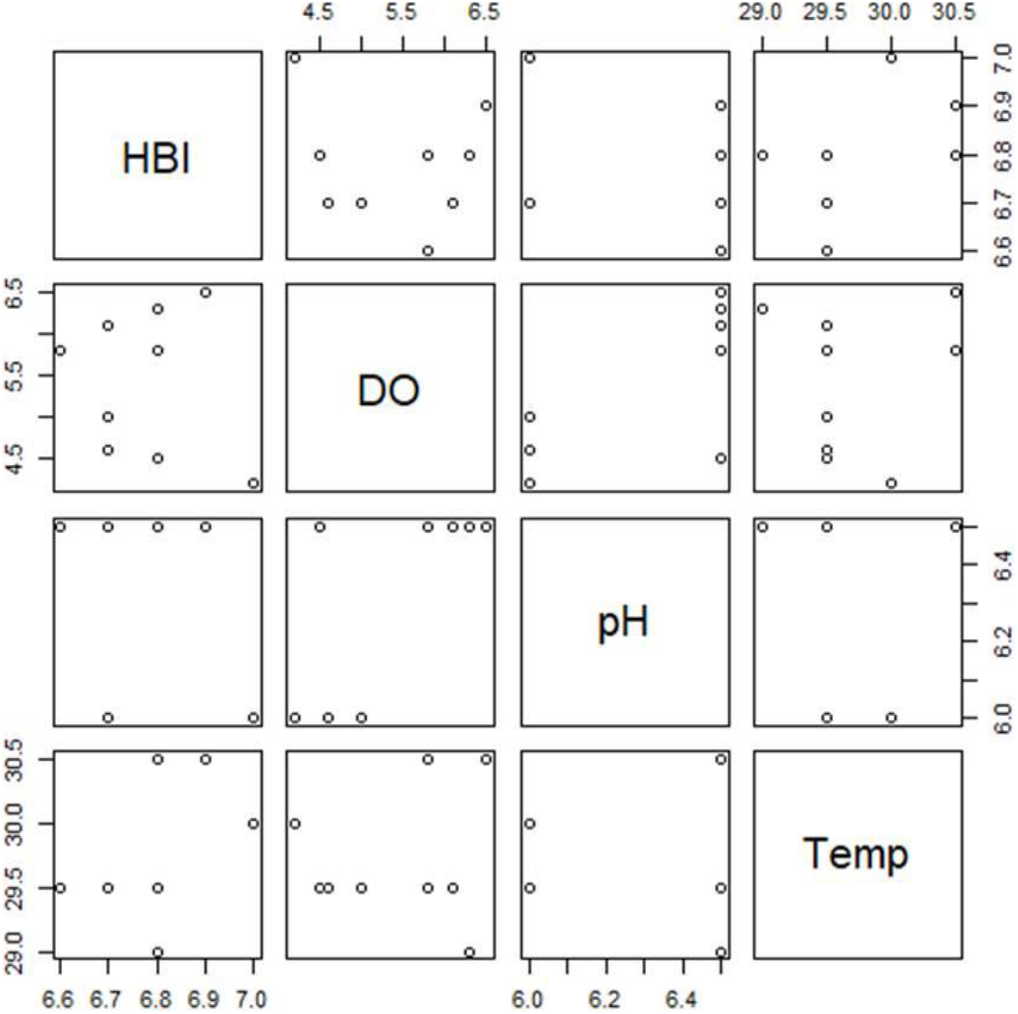
Correlations of HBI with water DO, pH, and temperature variables

### HBI and water quality variables

HBI shows varied correlation with DO, pH, and temperature. HBI has negative correlations with DO and pH whereas HBI has inverse correlation with temperature (Figure 4). Increase in DO has led to the decrease in HBI or increase in water quality since lower the HBI values then higher the water quality. For temperature variables, when the water becomes warmer, it is followed by an increase in HBI and a decrease in water quality.

**Figure 4.**
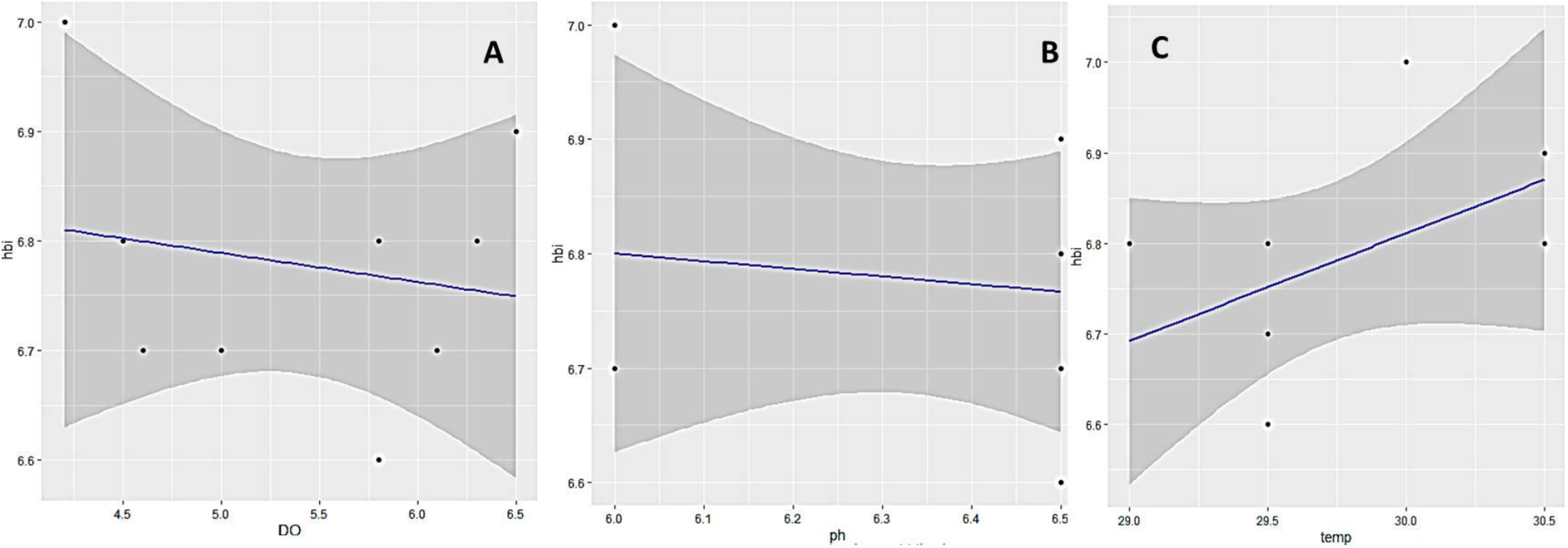
Linear regression of HBI with water DO (A), HBI with water pH (B), and HBI with water temperature

### HBI model selection

Akaike model selection confirms that the HBI and macrozoobenthos diversity can be used as a proxy representing water quality variables (Table 3). HBI model is the best to represent increase in water temperature (Ψ)HBI (~temp)(AIC = −10.264) (Figure 5) and also to represent water temperature increase and decrease in DO(Ψ)HBI (~temp+DO)(AIC = - 9.042398). Whereas HBI cannot be used to represent water pH (Ψ)HBI (~pH)(AIC = −7.830925).

**Table 3.** Akaike model selections for HBI, water DO, pH, and temperature variables

| Model | AIC | R<sup>2</sup> | Adj R<sup>2</sup> |
| --- | --- | --- | --- |
| ($\Psi$)<sub>HBI</sub> (~temp) | -10.26403 | 0.2516 | 0.1446 |
| ($\Psi$)<sub>HBI</sub> (~pH) | -7.830925 | 0.01923 | -0.1209 |
| ($\Psi$)<sub>HBI</sub> (~DO) | -7.982773 | 0.03564 | -0.1021 |
| ($\Psi$)<sub>HBI</sub> (~temp+pH) | -8.664654 | 0.2841 | 0.04553 |
| ($\Psi$)<sub>HBI</sub> (~temp+DO) | -9.042398 | 0.3136 | 0.08476 |
| ($\Psi$)<sub>HBI</sub> (~DO+pH) | -5.982928 | 0.03566 | -0.2858 |

**Figure 5.**
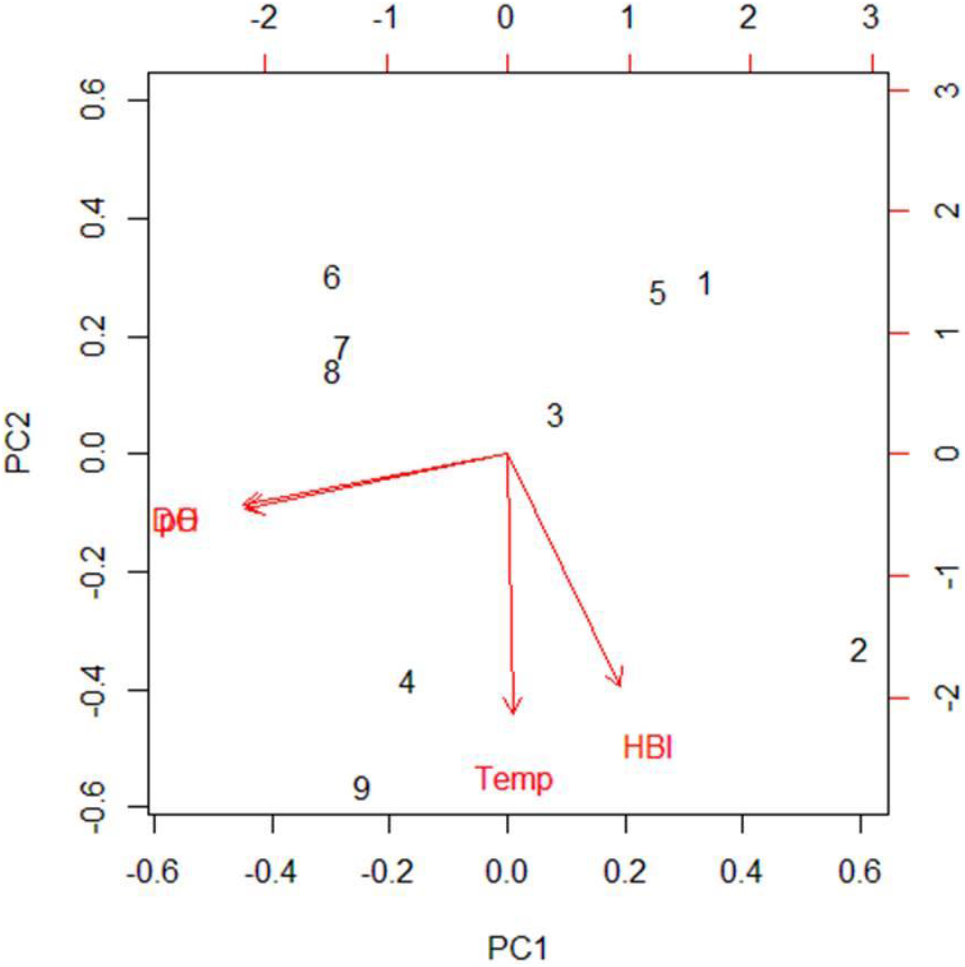
Principal Component Analysis of HBI with water DO, pH, and temperature

#### Discussion

In comparison to the current water quality assessments, the spatial HBI model in this study presents a novel method for describing precisely the distribution of HBI values in the space to allow accurate determination of the possible pollutant sources responsible for the higher HBI and decrease in water quality in lake. Analyzing these spatial distributions and patterns of HBI values in lakes using spatial model render them quantifiable and comparable. The output is very useful to evaluate the trends of water quality along land uses surrounding the lake. Besides that, the versatility of spatial model to visualize HBI is endorsing a broader application throughout the region or similar geographic areas. It possesses several intrinsic advantages worth noting.

The results obtained in this study are comparable to other studies (Amizera et al 2015, Du et al 2017) that are also using index to assess the water quality. A water quality variable that contributes more to the HBI values in this study is water temperature as observed by the model. In comparison to the research by Zamroni et al (2019), HBI in urban lake in this study is almost 3 fold higher than HBI in intact lake. This significant difference is related to the water temperature variables. With HBI equals to 6.6, the water temperature in urban lake in this study was higher than 29.5 °C. Whereas when the recorded water temperature was below 29.5 °C, HBI in intact lake is 2.2 and lower than HBI in urban lake.

Despite its versatility, HBI should be used with cautious. In classifying water quality at intact location, the HBI index may provide scores as expected. The HBI score is closed to the real time condition representing an ideal well-preserved environmental protection area condition with its intact riparian vegetation. This condition is related to the inlet condition of urban lake in this study where the obtained HBI values were low indicating good water quality. Whereas, Gonçalves & Menezes (2011) noticed that the index did not successfully reflect the obvious anthropogenic impacts there, since it is giving scores similar to those at intact locations. The explanation for this is considering that this index was developed originally for temperate systems and has not yet been adapted for tropical aquatic ecosystems. This reduces its usefulness for tropical environments since the index excludes a significant number of families found in tropics.

#### Conclusion

This study has confirmed 2 important matters. First, the water quality in the central parts of the urban lake was categorized as poor as indicated by high HBI. Second, the HBI is best to represent the increase in temperature variables and least feasible to predict pH variables.

